# The fitness costs and benefits of trisomy of each *Candida albicans* chromosome

**DOI:** 10.1101/2021.01.21.427689

**Authors:** Feng Yang, Yuan-ying Jiang, Yong-bing Cao, Judith Berman

## Abstract

*Candida albicans* is a prevalent human fungal pathogen. Rapid genomic change, due to aneuploidy, is a common mechanism that facilitates survival from multiple types of stresses including the few classes of available antifungal drugs. The stress survival of aneuploids occurs despite the fitness costs attributed to most aneuploids growing under idealized lab conditions. Systematic study of the aneuploid state in *C. albicans* has been hindered by the lack of a comprehensive collection of aneuploid strains. Here, we describe a collection of diploid *C. albicans* aneuploid strains, each carrying one extra copy of each chromosome, all from the same genetic background. We tested the fitness of this collection under several physiological conditions including shifts in pH, low glucose, oxidative stress, temperature, high osmolarity, membrane stress and cell wall stress. We found that, most aneuploids, under most conditions, were less fit than their euploid parent, yet there were specific conditions under which specific aneuploid isolates provided a fitness benefit relative to the euploid parent strain. Importantly, this fitness benefit was attributable to the change in the copy number of specific chromosomes. Thus, *C. albicans* can tolerate aneuploidy of each chromosome and some aneuploids confer improved growth under conditions that the yeast encounters in its host niches.

---

*Candida albicans* is the most prevalent human fungal pathogen (Pfaller *et al.* 2019). Aneuploidy, an unbalanced genomic state in which whole or segmental chromosomes, are missing or supernumerary, is often detected in yeasts and can be important for adaptation to stress conditions (reviewed in (Tsai and Nelliat 2019)). The first reported aneuploid karyotype in *C. albicans* that promoted stress adaptation was chromosome 5 monosomy (Chr5×1), which facilitated growth on L-sorbose as the sole carbon source (Janbon *et al.* 1998). In addition to whole chromosome aneuploidies, segmental aneuploidies alone can confer beneficial phenotypes under stress conditions. A classic example is isochromosome 5L (i(5L)), which was acquired recurrently in clinical azole-resistant *C. albicans* isolates (Selmecki *et al.* 2006). i(5L) confers azole resistance because it provides two additional copies of two genes: *ERG11,* which encodes lanosterol 14-alpha-demethylase, an enzyme critical for ergosterol biosynthesis and the target of azole drugs, and *TAC1,* which encodes a positive regulator of efflux pump gene expression (Selmecki *et al.* 2008). Aneuploidy also can affect responses to other antifungal drugs. For example, chromosome 2 trisomy (Chr2×3) and Chr5×1 promote adaptation to caspofungin (Yang *et al.* 2017; Yang *et al.* 2019).

In *C. albicans*, certain chromosome changes recurrently appeared under specific drug selection conditions (Selmecki *et al.* 2008; Yang *et al.* 2013; Ford *et al.* 2015; Yang *et al.* 2017; Yang *et al.* 2019), while others did not. Karyotype-specific properties also appear in *S. cerevisiae*: aneuploid strains with similar karyotypes have similar patterns with respect to their ability to grow across a range of different conditions (Pavelka *et al.* 2010). Different aneuploidies clearly give rise to different phenotypes, presumably because of specific genes that are present in altered dosage on the aneuploid chromosome (Selmecki *et al.* 2008; Sionov *et al.* 2010; Will *et al.* 2010). Furthermore, some aneuploidies yield multiple phenotypes that are attributable to different sets of gene(s) on the same aneuploid chromosome (Yang *et al.* 2019). Some phenotypes may be associated with the aneuploid state *per se*. In *S. cerevisiae,* many aneuploid strains are sensitive to conditions that interfere with protein synthesis and protein folding, which require more energy (Torres *et al.* 2007). Furthermore, aneuploids exhibit general hypo-osmotic-like stress and endocytic defects and a dependence on the Art-Rsp5 pathway, due to proteome imbalance (Tsai *et al.* 2019). The ability to distinguish between phenotypes that are chromosome-specific and those that are general properties of aneuploidy ideally requires examination of all possible aneuploids. Thus, a systematic collection of *C. albicans* aneuploid strains would provide an important tool for distinguishing chromosome-specific phenotypes from general aneuploid properties.

In *S. cerevisiae*, two collections of disomic haploid stains were isolated using genetic manipulations (Torres *et al.* 2007; Pavelka *et al.* 2010). Because *C. albicans* does not appear to undergo meiosis, different approaches to obtain *C. albicans* aneuploids are required. This includes using the parasexual mating cycle, in which diploid cells of opposite mating type first mate to form tetraploid cells, and then undergo concerted chromosome loss to return to the diploid or near-diploid state (Bennett and Johnson 2003), which often results in trisomic progeny, despite the lack of meiosis.

Given that many aneuploid chromosome combinations have been observed in *C. albicans*, it has long been suggested that *C. albicans* is more tolerant of aneuploidy than *S. cerevisiae* (Bouchonville *et al.* 2009). Aneuploidy clearly affects cell fitness differently under different growth conditions. In lab experiments in rich medium, *S. cerevisiae* disomic derivatives of haploids have fitness defects while trisomic derivatives of diploid isolates, as well as wild strains often exhibit lower fitness costs (Torres *et al.* 2007; Pavelka *et al.* 2010; Will *et al.* 2010; Filteau *et al.* 2015; Peter *et al.* 2018; Hose *et al.* 2020). Here we used a collection of *C. albicans* aneuploids to test this hypothesis for different trisomic homologs of each chromosome. We found that, in standard lab medium, all trisomic *C. albicans* strains were less fit than the diploid parent, and tetrasomic isolates as well as monosomy resulted in much lower fitness levels. Yet, specific aneuploidies conferred more rapid growth or higher biomass accumulation under some physiological stress conditions. This highlights the utility of this collection of strains, which has the potential to identify a broad range of phenotypes that may be altered by chromosome-specific and/or general features of aneuploidy.

## Results and discussion

We used SC5314, a diploid laboratory strain with 8 pairs of highly heterozygous chromosomes and an established haplotype (Van HET HOOG *et al.* 2007; Legrand *et al.* 2008; Muzzey *et al.* 2013). Over the years, we identified aneuploid isolates using a wide range of different selection pressures such as antifungal drug caspofungin, and chemotherapeutic drug hydroxyurea (Yang *et al.* 2017; Yang *et al.* 2019). In addition, we also screened for mutants able to survive from azoles (fluconazole, ketoconazole, miconazole), another two members of echinocandins (micafungin, anidulafungin), flucytosine, amphotericin B, sphingolipid pathway inhibitors (myriocin and aureobasidin A), endoplasmic reticulum stress inducers (tunicamycin and brefeldin A), and other chemotherapeutic drugs such as fluorouracil (Yang et al, manuscript in preparation). Combining these collections allowed us to collect many aneuploid isolates, all derived from lab strain SC5314. The collection includes trisomic isolates with extra copies of each homolog of each chromosome (AAB and ABB), isolates with chromosomes 2 or 5 tetrasomy and chromosome 5 monosomy (Fig 1).

**Figure 1.**
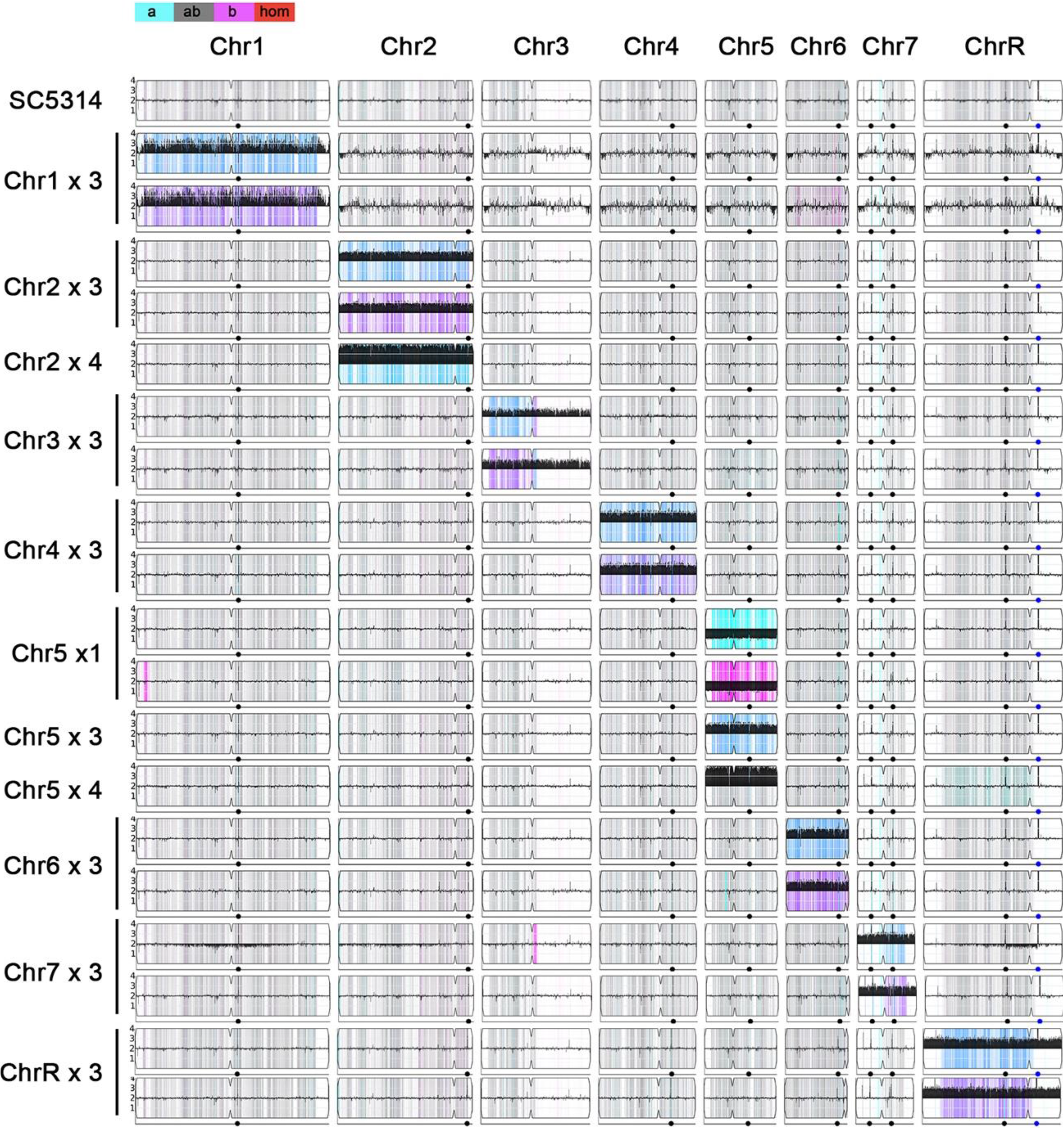
Ymap karyotypes of *C. albicans* aneuploid collection. Representative karyotypes of singly trisomic, tetrasomic and monosomic strains derived from parent strain SC5314 generated using Ymap (Abbey *et al.* 2014) as described in (Yang *et al.* 2019). Read depth was normalized to that of the diploid parent and is shown on the y-axis on a log_2_ scale converted to absolute copy numbers (1-4). Allelic ratios (A:B) are color-coded: grey, 1:1 (A/B); cyan, 1:0 (A or A/A); magenta, 0:1 (B or B/B); purple, 1:2 (A/B/B); blue, 2:1 (A/A/B); light blue, 3:1 (A/A/A/B)).

Aneuploid isolates were initially identified based upon three criteria for phenotypic variation: first, improved growth, relative to the parent, in the presence of (at least one) specific stress; second, instability of the acquired adaptive state when selection was removed, which was usually detected as the formation of a mixture of smaller (S) and larger (L) colonies on non-selective medium (Yang *et al.* 2019) (Fig. 2A); and third, loss of the adaptive stated from the L colonies taken from non-selective medium (Yang *et al.* 2019). Next-generation sequencing of colony progeny that fulfilled the three criteria revealed aneuploidy in most isolates. Furthermore, the L derivatives of the initially aneuploid isolates (those that had lost the adaptive state) were euploid, consistent with the idea that the aneuploid chromosome facilitated growth in the presence of the relevant drug and that loss of the aneuploidy, in the absence of selection, was accompanied by the loss of the adaptive phenotype (Yang *et al.* 2013; Yang *et al.* 2019). To test the fitness of the aneuploid collection, we first monitored growth dynamics in rich YPD medium. All the aneuploid strains (the S colonies) were less fit than their parent, while the L colonies usually grew with fitness comparable to the parent (Fig 2B). Of note, trisomy of many of the chromosomes often incurred only a small, or no, fitness cost, and only Chr2 showed substantively different growth rates between the two homologs. Isolates monosomic for Chr5 also grew very slowly, with the Chr5×1A homolog more fit than the Chr5×1B homolog. Similarly, Chr5×1 A was found more frequently as a caspofungin survivor than Chr5×1 B (Yang et al. 2019). Importantly, the spontaneous homozygous diploid derivatives of these monosomic strains formed large colonies and grew with kinetics like those of the parental strain. Thus, we suggest that the slower growth of the monosomic isolates is not due to lack of homolog-specific alleles, but rather due to copy number sensitivity of those alleles. Furthermore, Chr5×4 (1.1 MB) (AABB) had no obvious fitness cost, while Chr2×4 (~2.2 MB) reduced growth more than the Chr2×3 isolates. While chromosome size (and the accompanying additional burden of extra DNA) might contribute to this difference, this cannot be the sole reason, given that trisomy of Chr7 (950 kb, the smallest chromosome) had a similar fitness cost to trisomy of Chr1 (3.2 MB, the largest chromosome).

**Figure 2.**
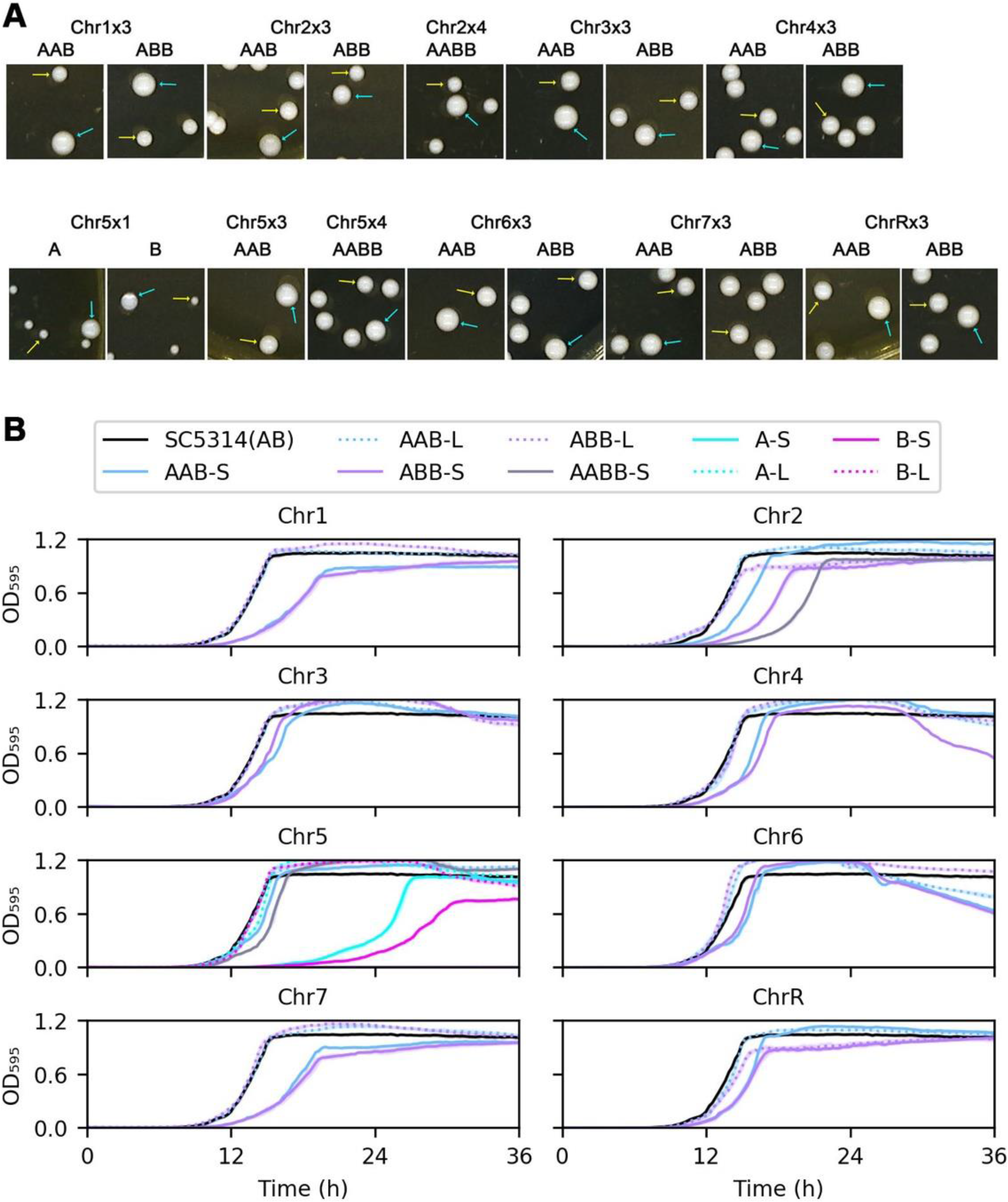
Effects of aneuploidies on fitness in rich medium. In A, approximately 200 colonies of each putative aneuploid strain were plated on YPD agar plates, incubated at 37’C for 36 h and then photographed. Yellow arrows indicate small (S) colonies. Cyan arrows indicate large (L) colonies. In B, one small colony (S) and one large colony (L) for each isolate were chosen from YPD plates. Growth was monitored every 15 min at 37’C for 36 h. Data are represented as the mean ± SD of three technical replicates.

We also analyzed the fitness of the aneuploid collection on several stress conditions that reduce the growth of euploid strains including environmental perturbations, such as low (5.0) and high pH (8.0), nutrient limitation (0.1% glucose), oxidative stress (hydrogen peroxide, 6 mM), elevated temperature (39°C, as a proxy for febrile temperature), and high osmolarity (1.2M NaCl) (Fig. 3). We found a wide range of phenotypic variation across the aneuploid strains, and under particular stress condition, particular aneuploidy conferred better fitness than the euploid parent. Chr7×3 (ABB) was better fit in the presence of H_2_O_2_, Chr3×3 (AAB and ABB), Chr6×3 (AAB) and Chr7×3 (ABB) were better fit at low pH. Under other stress conditions, the fitness difference between most trisomy strains and the euploid parent was also not as obvious as the difference when grown in the absence of stress.

**Figure 3.**
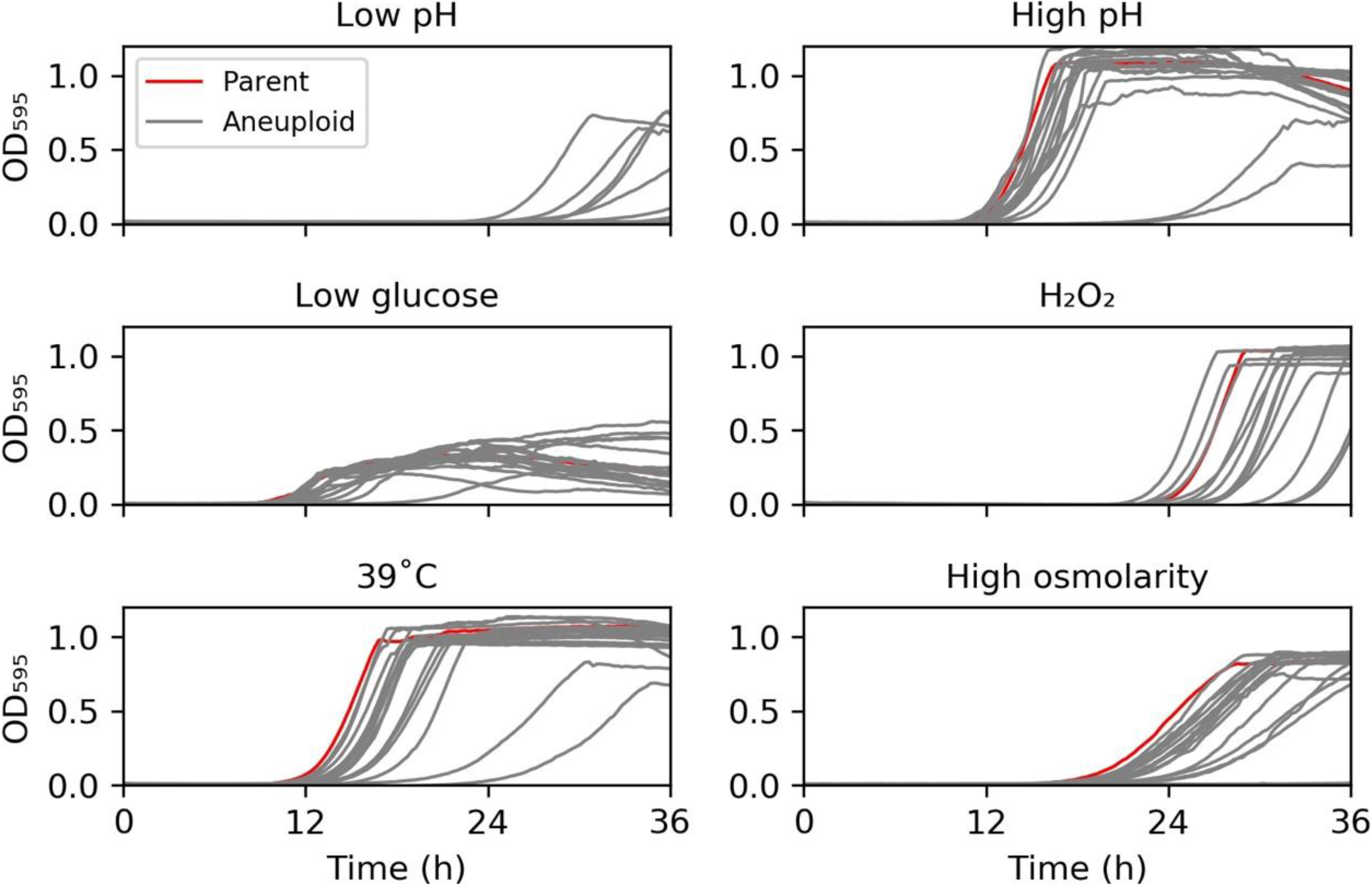
Fitness under physiological stress conditions. SC5314 and the aneuploid collection were grown in YPD medium in 96 well plates under the following conditions: low pH (YEPD adjusted to pH 5.0), high pH (YPD with pH 8.0), low glucose (YPD with 0.1% glucose), oxidative stress (YPD supplemented with 6 mM H_2_O_2_), febrile temperature (39°C). and high osmolarity (YPD with 1.2 M NaCl). The plates were incubated at 37°C unless otherwise specified. Data represent the mean ± SD of three technical replicates.

To analyze fitness during exposure to cell membrane and cell wall stresses, we exposed the strains to sodium dodecyl sulfate (SDS) and caffeine. SDS is a detergent that disrupts cell membranes, activates stress responses including the cell wall integrity (CWI) pathway, and restricts cell growth (Kono *et al.* 2012); caffeine elicits pleiotropic effects including inhibiting cell growth, reducing cell fitness, arresting the cell cycle, and ultimately leads to cell death (Kuranda *et al.* 2006)(Fig. 4A). Interestingly, Chr6×3 (both AAB and ABB) grew better than the diploid parent strain on both SDS and caffeine. Given that Chr6×3 in an SC5314 derivative exhibited reduced virulence and increased commensal growth ability (Forche *et al.* 2019), we suggest that higher tolerance to cell membrane and cell wall stresses may underpin the improved ability of Chr6×3 trisomic isolates to colonize and reside in the host gastrointestinal tract.

**Figure 4.**
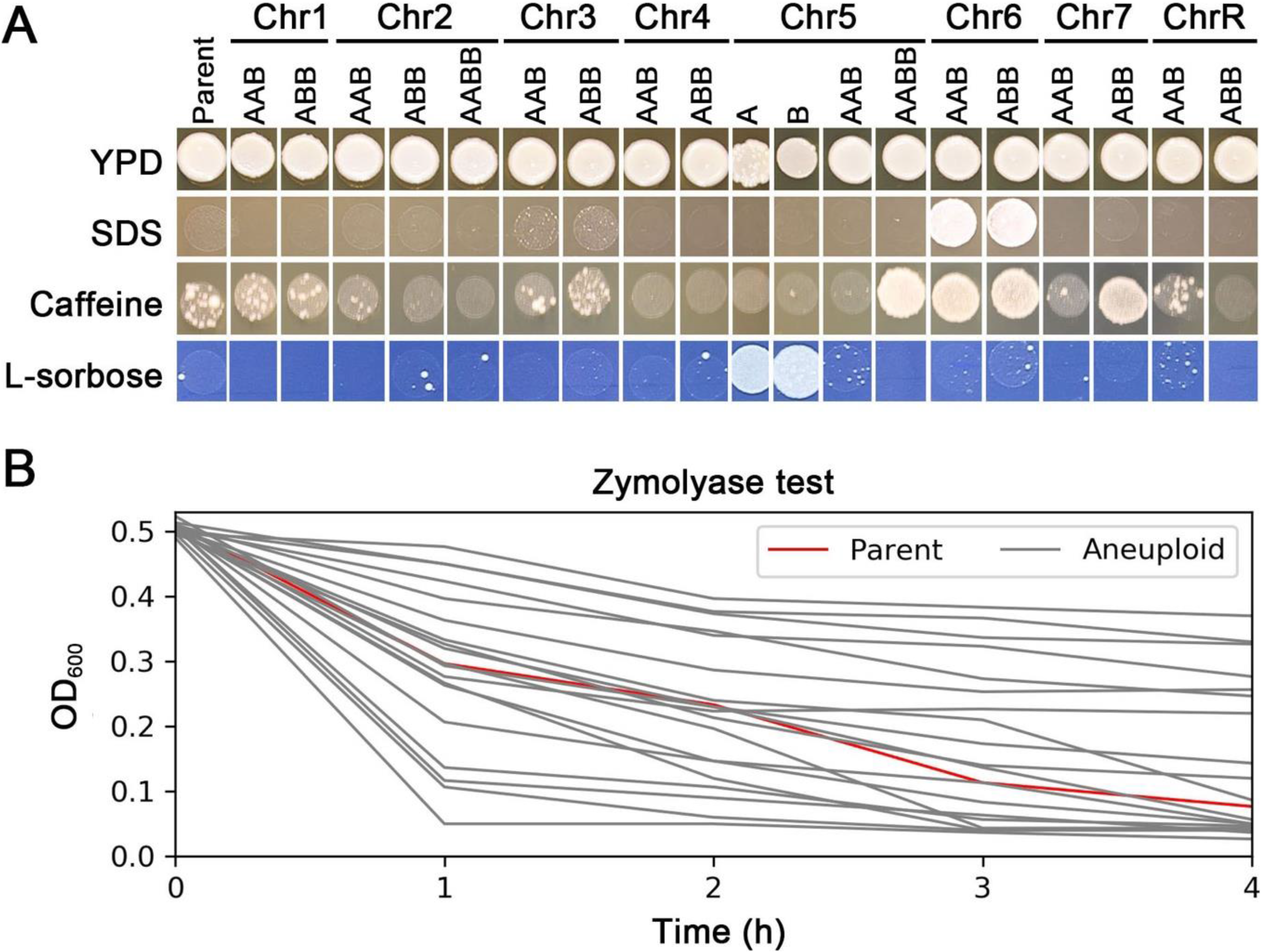
Effects of aneuploidies on membrane stress and cell wall stress. In A, spot assays identify aneuploidies that confer increased and decreased fitness in different stresses. Comparison of parent and all trisomic, monosomic and tetrasomic aneuploid strains for the ability to grow on sodium dodecyl sulfate (SDS) or Caffeine, or sorbose (6.7 g/L yeast nitrogen base without amino acids, 2% L-sorbose, 2% agar). Each spot contained approximately 3.0×10^4^ cells. Plates were incubated at 37°C for 3 days and then photographed. In B, the parent (red line) and the aneuploid strains were suspended in 10 mM Tris-HCl (pH 7.5) containing 50 μg/ml of zymolyase. Cell suspensions were maintained at 37°C for 4 h. The optical density was measured at 1h time intervals. Each data point was the mean ± SD of three technical replicates.

**Figure 5.**
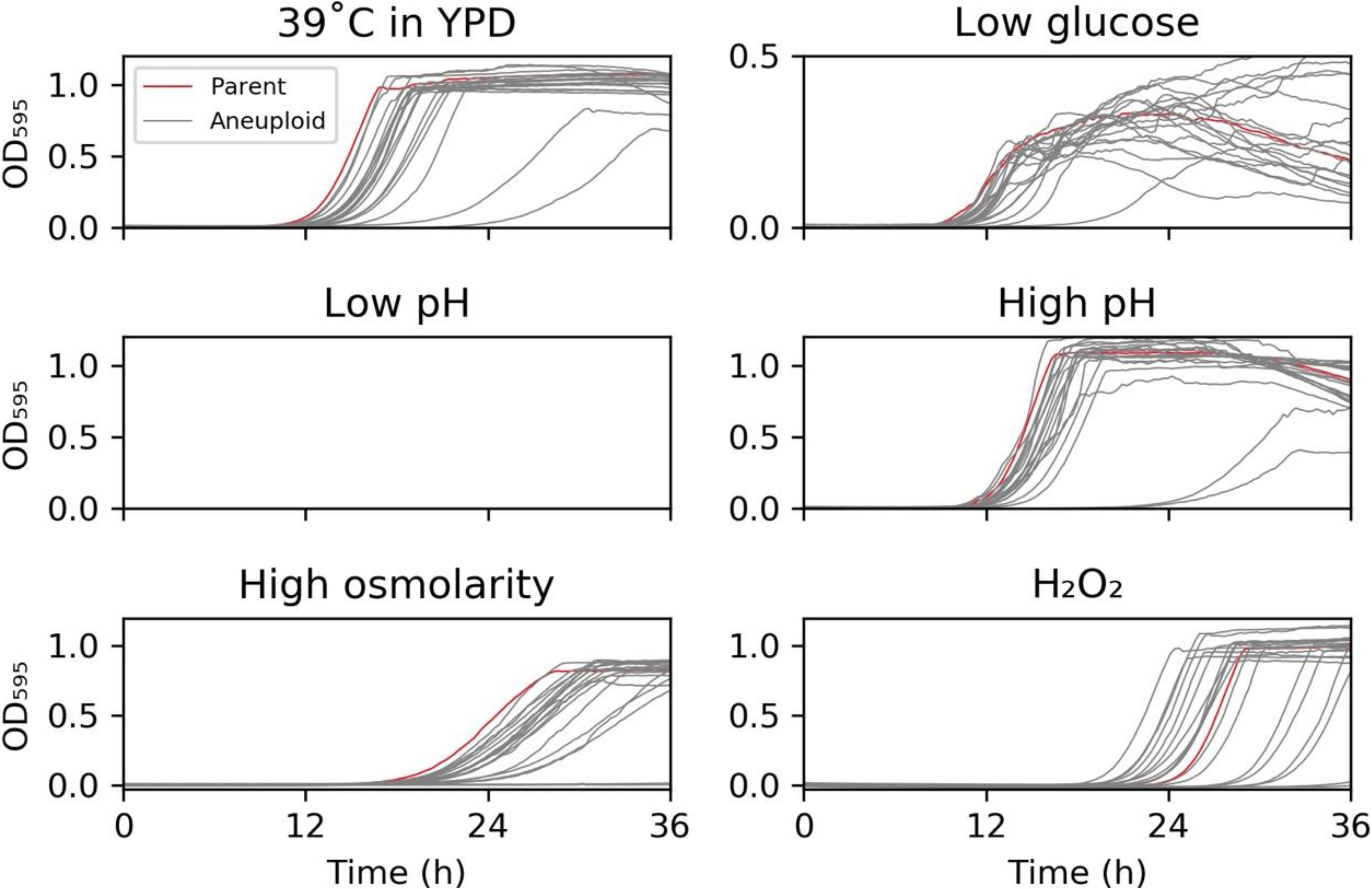

Relative to the euploid parent, several of the isolates, including Chr7×3 ABB, had a fitness benefit on caffeine while several other aneuploids, including Chr7 AAB had a fitness cost on caffeine. This suggests that the aneuploidy affects growth in caffeine stress in an allele-specific manner. Alternatively, the papillated growth seen in the euploid parent on caffeine may be due to the appearance of new aneuploidies that arise on the caffeine-containing plate, perhaps because caffeine can induce aneuploidy through asymmetric cell divisions (Katsuki *et al.* 2008). In this alternative model, specific genes on different aneuploid chromosomes likely affect both the ability to withstand the growth-inhibiting effects of caffeine, as well as the susceptibility to caffeine-enhanced genome instability. As expected (Janbon *et al.* 1998), Chr5×1, and no other aneuploid isolates grew well on sorbose. Finally, there was a wide range of sensitivity to zymolyase digestion, a lytic enzyme that digests the *C. albicans* cell wall (Fig. 4B).

In conclusion, *C. albicans* can tolerate trisomy of each chromosome. In the absence of stress, aneuploid strains are less fit, but can spontaneously revert to better fitness by reverting to euploidy. Under particular stress conditions, some aneuploid chromosomes confer better fitness than the euploid parent.

## Materials and Methods

### Identifying aneuploid isolates

*C. albicans* reference strain SC5314 was streaked from −80°C glycerol stocks to YPD agar and incubated at 37°C for 24 h. Several randomly chosen colonies were suspended in distilled water, cell density was determined using a Cellometer (Nexcelom), and approximately 1.0×10^6^ cells were spread onto YPD plates supplemented with drugs at concentrations that inhibited growth of the vast majority of SC5314 cells (Yang *et al.* 2017; Yang *et al.* 2019; Yang *et al.,* manuscript in preparation). The plates were incubated at 30°C (fluconazole) or 37°C (other drugs) for 3 days. Rare colonies that appeared were randomly chosen and saved in −80°C glycerol stocks.

### Colony instability assay

Aneuploid strains were streaked from −80°C freezer to YPD agar and incubated at 37°C for 36 h. One small colony was randomly chosen and suspended in distilled water. Cells were diluted with distilled water and approximately 200 cells were spread on a YPD plate and incubated at 37°C for 36 h. One small (S) colony and one large (L) colony were randomly chosen for further studies.

### Fitness assay

Approximately 1×10^3^ cells/ml were suspended in the medium specified in figure legends and 150 μl was transferred to 96-well plates. Optical density (OD_595nm_) was measured every 15 min for 36 h at 37°C using a Tecan plate reader (Infinite F200 PRO, Tecan, Switzerland).

### Spot assay

Strains were streaked onto YPD agar, incubated at 37°C for 36 h (aneuploid strains) or 24 h (parent) and several colonies were chosen randomly and suspended in distilled water. Cell densities were adjusted to 1×10^7^ cells/ml and 3 μl of 10-fold serial dilutions was spotted on YPD agar plates supplemented with the compounds described in figure legends. The plates were incubated at 37°C for 3 days and then photographed.

### Assay of sensitivity to zymolyase

Cells were suspended in 10 mM Tris-HCl (pH 7.5) containing 50 μg/ml of zymolyase 20T (U.S. Biological, Swampscott, MA). Cell density was adjusted to OD_600_ of 0.5, incubated at 37°C for 4 h and the decrease in optical density was monitored at 1h time intervals.

**Next generation sequencing** was performed as described previously (Yang *et al.* 2017).

## Data availability

The genome sequence data are available at via ArrayExpress at EMBL-EBI (www.ebi.ac.uk/arrayexpress) under accession number E-MTAB-9739.

## Acknowledgments

This project was supported by National Natural Science Foundation of China (No. 82020108032 to YYJ, No. 81673478, and No. 81872910 to YBC), Shanghai Key Basic Research Project (No. 19JC1414900 to YB C), and the Israel Science foundation (#997/18 to JB). We apologize to authors whose work has not been cited due to limitations of space. We thank Dr. Hung-Ji Tsai for critical reading of the manuscript.

